# Single-cell imaging reveals non-cooperative and cooperative infection strategies of *Listeria monocytogenes* in macrophages

**DOI:** 10.1101/2022.06.04.493993

**Authors:** Josephine Moran, Liam Feltham, James Bagnall, Marie Goldrick, Elizabeth Lord, Catherine Nettleton, David G. Spiller, Ian Roberts, Pawel Paszek

## Abstract

Pathogens have developed intricate strategies to overcome the host’s innate immune responses. In this paper we use live-cell microscopy with a single bacterium resolution to follow in real time interactions between the food-borne pathogen *L. monocytogenes* and host macrophages, a key event controlling the infection *in vivo*. We demonstrate that infection results in heterogeneous outcomes, with only a subset of bacteria able to establish a replicative invasion of macrophages. The fate of individual bacteria in the same host cell was independent from each other and non-cooperative, but a higher multiplicity of infection resulted in a reduced probability of replication. Using internalisation assays and conditional probabilities to mathematically describe the multi-stage invasion process, we demonstrate that the secreted Listeriolysin toxin (LLO) of the PrfA regulon regulates replication probability by compromising the ability to phagocytose bacteria. Using strains expressing fluorescent reporters to follow transcription of either the LLO-encoding *hly* or *actA* genes, we show that replicative bacteria exhibited higher PrfA regulon expression in comparison to those bacteria that did not replicate, however elevated PrfA expression *per se* was not sufficient to increase the probability of replication. Overall, this demonstrates a new role for the population-level, but not single cell PrfA-mediated cooperativity to regulate outcomes of host pathogen interactions.

**Key points:**

- *L. monocytogenes* invasion of innate immune macrophages results in heterogeneous infection outcomes at the single cell level
- Fate of individual bacteria in the same host cell is independent from each other and non-cooperative
- Bacterial populations coordinate host cell uptake via the rate of phagocytosis to reduce internalization at high MOI
- The PrfA regulon system is necessary but not sufficient for *L. monocytogenes* replication, but population-level PrfA virulence regulates single cell outcome probability

## Introduction

Specific interactions between pathogenic bacteria and individual host cells decide the course of an infection and its’ outcome. The responses of individual host cells are extremely variable, as exhibited by noisy transcription factor dynamics (Adamson et al. 2016; Czerkies et al. 2018; Kellogg et al. 2017; Cheng et al. 2015; Patel et al. 2021) and heterogeneous effector gene production (Shalek et al. 2014; Bagnall et al. 2018; Bagnall et al. 2020; Xue et al. 2015). In turn, pathogens employ complex strategies to avoid recognition by host cells (Rosenberger and Finlay 2003; Nikitas et al. 2011; Avraham et al. 2015), and are able to rapidly adapt to environmental changes to diversify their phenotypes and enhance their survival in the host (Norman et al. 2015). Consequently, the interactions between host and pathogen at the single cell level are inherently heterogeneous and result in different and “seemingly” probabilistic outcomes (García-Del Portillo 2008; Helaine et al. 2010). For example, only a subset of genetically identical host cells can kill invading *Salmonella* (McIntrye, Rowley, and Jenkin 1967), while others allow a pathogen to either persist or replicate to eventually cause a systemic infection (Avraham et al. 2015; Stapels et al. 2018). Whether different infection outcomes are controlled by the pathogen, the host, or both is not well understood.

Here we use real time single cell analyses to study the food-borne pathogen *Listeria monocytogenes* which is responsible for a number of serious infections with high mortality rates (20-30% in human) despite antibiotic intervention (Cassir, Rolain, and Brouqui 2014). The potential of *L. monocytogenes* to cause systemic infection depends on the ability to transcytose the intestinal barrier and subvert immune cells to establish infections in the liver and spleen (Nikitas et al. 2011; Kim et al. 2021). *L. monocytogenes* invades host cells via a membrane-bound vacuole (through phagocytosis by immune cells), before escaping, replicating in the cytoplasm, and spreading to adjacent cells, coordinated through the action of the regulatory protein PrfA (Radoshevich and Cossart 2018). The PrfA regulon contains genes required for invasion of non-phagocytic cells, phagosome escape (*hly* encoding pore-forming toxin listeriolysin O, LLO), cytosolic growth and spread to neighbouring cells through actin polymerisation (*actA*) (Cossart 2011; Wang et al. 2015). Regulation of PrfA activity is complex, involving transcriptional and posttranslational control (Radoshevich and Cossart 2018; Reniere et al. 2015; Johansson et al. 2002; Krypotou et al. 2019). Recently, it has been shown that the response of *L. monocytogenes* at the single cell level to environmental triggers was heterogeneous, where only a subset of *L. monocytogenes* expressed the PrfA regulated *hly* (Guldimann et al. 2017). Likewise, in epithelial cells a small sub-population of pioneer *L. monocytogenes* promoted enhanced cell-to-cell spread (Ortega, Koslover, and Theriot 2019). *L. monocytogenes* is also capable of switching between different phenotypic states inside the host to diversify its invasion strategies, from an active motile to persistent non-replicative state (Kortebi et al. 2017). In turn, genetically identical host cells exhibit different susceptibility to *L. monocytogenes* invasion through the heterogeneity of the endothelial cell adhesions (Rengarajan and Theriot 2020). Despite the recent advances highlighting the heterogeneous nature of interactions between bacterial pathogens and host cells, our mechanistic understanding how the variability in the pathogen and in the host contribute to the overall outcome of infection at the single cell level is limited.

Here we use live-cell confocal microscopy approaches with single bacteria resolution to understand interactions between *L. monocytogenes* and host macrophages, a critical event controlling infection (Shaughnessy and Swanson 2007). We show that infection relies on a fraction of bacteria that can effectively replicate and spread within the macrophage population. We demonstrate that the ability of *L. monocytogenes* to replicate is non-cooperative as multiple bacteria in the same host cell have statistically independent fates, but the overall probability is controlled by the multiplicity of infection (MOI). We demonstrate that this is regulated through the secreted LLO in the environment, which compromises macrophages’ ability to phagocytose *L. monocytogenes* at higher MOI. Paradoxically, we found that at the single cell level PrfA regulon expression is heterogeneous and positively correlates with infection outcomes, however it is not sufficient for *L. monocytogenes* replication. Overall, these data provide new insights into PrfA-mediated interactions of *L. monocytogenes* and innate immune macrophages and potential new avenues to manipulate infection outcomes at the single cell level.

## Results

### Infection of macrophages results in heterogeneous outcomes at the single cell level

To quantify outcomes of individual interactions of *L. monocytogenes* and host macrophages we used live cell confocal microscopy approaches. We infected monolayers of a murine macrophage cell line, (RAW 264.7) and primary murine bone marrow derived macrophages (BMDMs) with *L. monocytogenes* expressing green fluorescent protein (referred herein as *Lm*-GFP) using a chromosomally integrated plasmid system (see Materials and Methods). We used a gentamycin protection assay (where start of the imaging experiment in referred as t0) and a low MOI of 0.25 (4:1 host cell to pathogen ratio) to exclude multiple invasion events per host cell and thus spatially separate individual host-pathogen interactions (Fig. 1a). In a typical experiment this resulted in 4.1% of host cells (90 per 2200 cells) harbouring exactly 1 bacterium at t0 (with 1% of infected host cells harbouring >1 bacteria). In contrary to previous approaches, which typically quantify the “average” behaviour of many bacteria per host, our approach therefore enables analyses of individual host cell pathogen interactions with a single bacterium resolution in a more physiological and clinical context.

**Figure 1.**
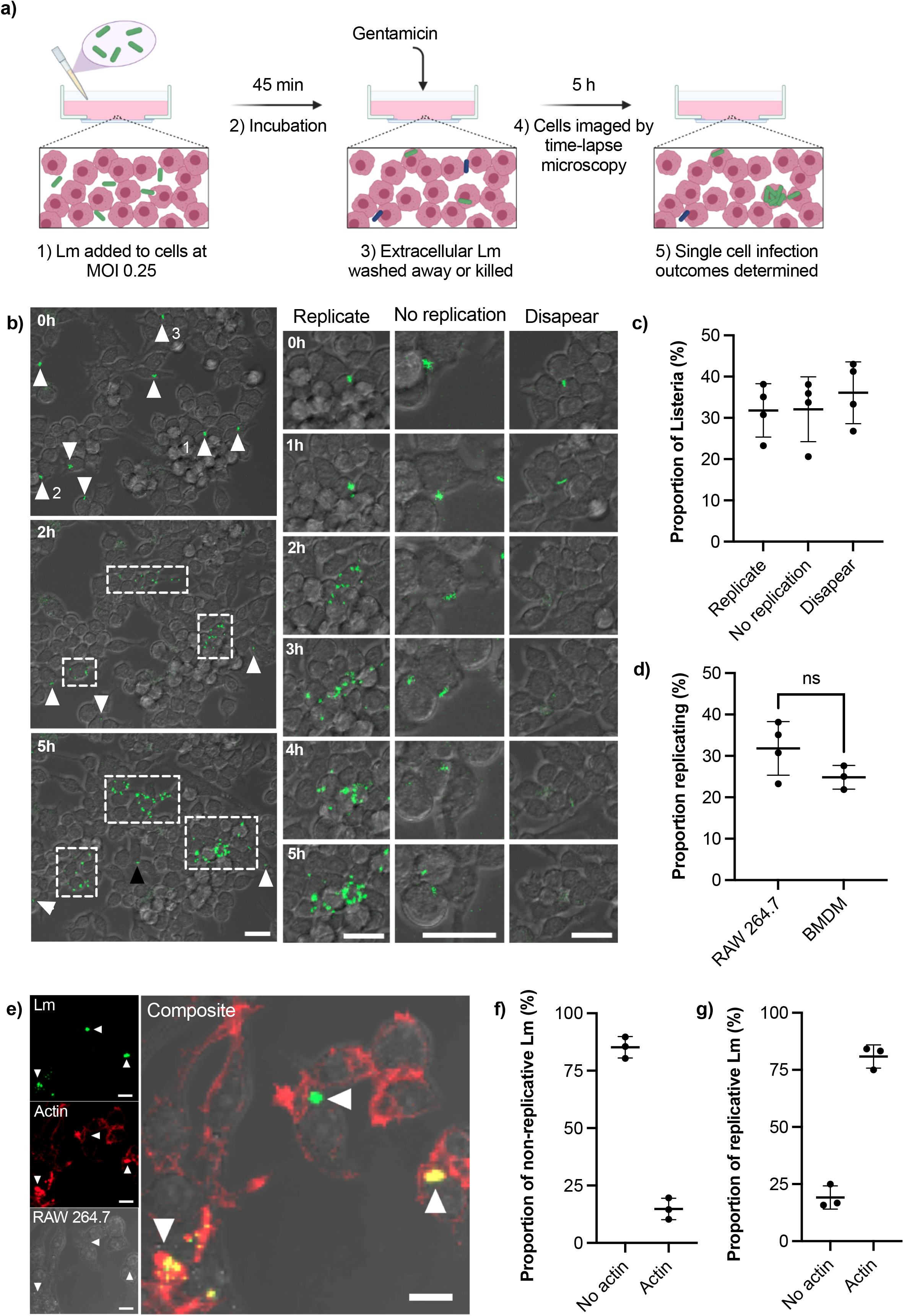
Heterogeneous outcomes of *L. monocytogenes* infection of macrophages at the single cell level. a) Schematic representation of infection protocol: 1) RAW 264.7 macrophages infected with *Lm*-GFP at an MOI of 0.25; 2) Cells and *Lm*-GFP are incubated for 45 mins; 3) Non-adherent *Lm*-GFP are washed away and fresh media containing gentamycin is added to inhibit growth of extracellular bacteria (referred herein as t0); 4) Sample is imaged by time-lapse confocal microscopy for 5 h to determine infection outcomes. b) Representative live-cell microscopy images of RAW 264.7 macrophages (brightfield) infected with *Lm*-GFP (green) from times 0-5 h post gentamycin treatment. White arrows-non-replicative *Lm*-GFP present at t0; white boxes-replicative foci and black arrow: single *Lm*-GFP not visible at t0. The right panels show magnified examples of the 3 outcomes resulting in: (1) replicative infection; (2) non-replicative infection and (3) disappearance. Scale 20 μM. c) Proportion of different single cell infection outcomes as depicted in b evaluated at 5 h as a function of the total *Lm*-GFP interactions at t0. Data from four replicates (from 358 individual interactions) shown in circles with mean and SD as solid lines. d) Proportion of replicative invasions for primary BMDMs infected with *Lm*-GFP, evaluated at 5 h as a function of the total *Lm*-GFP associated with BMDMs at t0 (in comparison to data from c). Triplicate data (from 153 individual interactions) shown in circles with mean and SD as solid lines. Statistical significance (ns = non-significant) assessed using Mann-Whitney rank test. e) Representative image of actin staining showing: *Lm*-GFP replicating (arrow pointing down), non-replicating without acting association (arrow pointing left) and non-replicating associated with actin (arrow pointing up). RAW 264.7 macrophages (brightfield) infected with *Lm*-GFP were fixed at 5 h, permeabilised then stained with anti-Lm (green) and phalloidin-594 (red). Scale bar 10 μM. Images representative of three replicated experiments. f) Proportion of non-replicative *Lm*-GFP at 5 h with or without actin staining colocalization as depicted in e. Triplicate data shown in circles with mean and SD as solid lines. g) Proportion of replicative *Lm*-GFP at 5 h with or without actin staining colocalization based on data in e. Triplicate data shown in circles with mean and SD as solid lines.

Upon infection (Fig. 1b, video 1) we identified three main outcomes at 5 h post infection resulting in: (1) intracellular replication of bacteria; (2) non-replicative invasion and (3) disappearance. We found that upon invasion of RAW 264.7 macrophages, on average only 32% (±6% standard deviation, SD) of individual host-pathogen interactions resulted in a replicative infection (Fig. 1c). These replication events were typically initiated within the 2 h post infection and resulted in a rapid growth (up to 30 bacteria in 5h) and spread to neighbouring host cells forming characteristic replicative foci (as indicated by white boxes in Fig. 1b). Likewise, 32% (±8%) of single-cell interactions resulted in non-replicative invasion, where individual bacteria remained associated with the original host cell for the duration of the experiment. In some cases, bacteria established a new interaction event at later time point, see black arrow in Fig. 1b. In addition, 36% (±8%) of bacteria disappeared within the 5 h post infection from the imaging region. The latter is consistent with phagosome killing of bacteria; however, we cannot exclude a possibility that some bacteria escaped to the media. Importantly, the invasion of murine BMDMs at MOI=0.25 resulted in similar infection outcomes; 25% (±3%) of interactions resulted in replicative invasion, which was not statistically different from the macrophage cell line (Fig. 1d),

The intracellular life cycle of *L. monocytogenes* is well characterised, to grow the bacteria must escape the phagosome to the cytoplasm, where it can replicate and accumulate actin for intracellular propulsion (Radoshevich and Cossart 2018). We therefore used phalloidin staining to assess the ability of *L. monocytogenes* to polymerase actin (Fig. 1e), as a marker of its cytoplasmic localisation (Kocks et al. 1992). We found that 15% (±6%) of non-replicative bacteria co-localised with actin staining (Fig. 1f) suggesting that these bacteria are present in the host cytoplasm, but do not replicate. The presence of actin would also indicate these bacteria were not destined for autophagy (Birmingham et al. 2007). In comparison, 81% (±5%) of replicative bacteria co-localised with actin (Fig. 1g). Overall, these data demonstrate the heterogeneous nature of interactions between *L. monocytogenes* and host macrophages with only a fraction of bacteria that can effectively replicate and spread within the host cell population.

### Invasion of individual bacteria in the same host cell is non-cooperative

The fundamental question that we wanted to address is whether the single cell infection outcomes are determined by the host or the pathogen, or both. To discriminate between these possibilities, we simultaneously infected macrophages with *L. monocytogenes* expressing either green (*Lm*-GFP) or red fluorescent protein (*Lm*-dsRed) using a combined MOI of 5 (at 1:1 ratio between red and green bacteria). This enabled us to determine outcomes of multiple invasion events per individual host cell. For example, if two bacteria share the same fate upon invasion of the same host cells (e.g., the ability to replicate) it would suggest that the single cell outcome is controlled by the host environment, i.e., some host cells are more permissive to replication than others. Conversely, if fates are statistically independent, the infection outcome is controlled by the bacteria (Fig. 2a).

**Figure 2.**
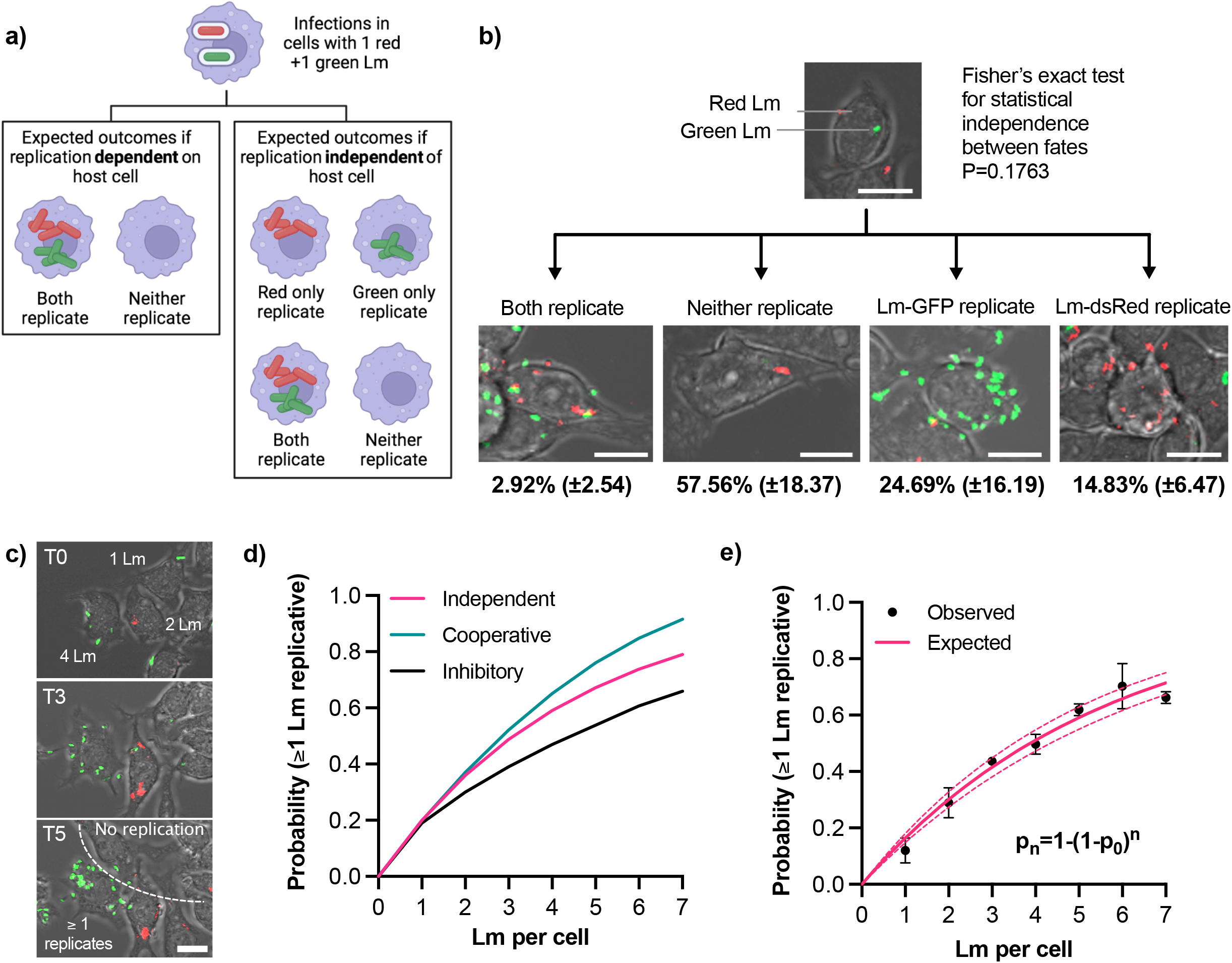
Fate of individual bacteria in the same host cell is independent. a) Experimental rationale: fates of red and green *L. monocytogenes* in the same host cell determine pathogen and host contribution to replicative invasion. If both bacteria share the same fate in the same host cell, the host environment controls the outcome, if fates are independent, bacteria control the outcome. b) Representative images of different outcomes of RAW 264.7 macrophages simultaneously infected with *Lm*-dsRed and *Lm*-GFP at combined MOI 5 (at 1:1 ratio). Shown are the mean proportion and SDs of different infection outcomes of triplicate data subset of cells with one *Lm*-dsRed and one *Lm*-GFP at t0 (total 167 cells). Scale bar 10 μM. c) Quantification of the replication probability for multiple invasion events per host cell. Representative images of data from b, with cells harbouring 1-4 bacteria at t0. Increased pathogen number over time (as highlighted on the image) indicates that at least one bacterium replicated. Scale bar 10 μM. d) Schematic representation of collective invasion strategies: (1) Cooperative invasion: multiple bacteria in the same cell promote each other’s replication, leading to increased replication probability; (2) non-cooperative invasion: bacterial replication is independent in the same host (probability of replication given by statistical independence p_n_=1-(1-p_0_)^n^, where n is the number of bacteria, p0 replication probability for 1 bacteria); (3) inhibitory invasion: reduced bacterial replication probability as number of bacteria increases due to enhanced immune response. e) Probability that at least one *L. monocytogenes* replicates as a function of number of bacteria per host cell at t0. Shown in black are observed probabilities (mean and SDs, based on three replicate experiments). Solid pink line depicts expected probabilities assuming statistical independence, and 95% confidence intervals in broken lines.

To test these possibilities, we first focused on a subset of host cells that at t0 were infected with exactly one green and red bacteria (167 cells from triplicate experiments, Fig. 2b, video 2). We found that the marginal probabilities that green or red bacteria replicate although slightly different from each other, p_G_= 0.257 (±0.16) vs. p_R_=0.148 (±0.07), respectively, were not statistically different (Mann-Whitney test p value 0.4). Assuming the statistical independence, the expected probability that both red and green bacteria replicate in the same host cell is the product of their marginal probabilities, p= p_G_ x p_R_ = 0.257 × 0.148= 0.036 (±0.04), while the expected probability that neither red nor green replicate is p_n_=(1-p_G_) x (1-p_R_)= 0.63 (±0.19). In the data we observed that the probability that both green and red bacteria replicated was 0.029 (±0.025), while probability that neither replicated was 0.58 (±0.18). These could not be statistically distinguished from the expected probabilities (Fisher exact test p value 0.18). Therefore, the fate of individual bacteria in the same host is independent from each other, suggesting it is the behaviour of individual *L. monocytogenes* cells that determine the overall outcome of the infection.

The presence of multiple bacteria per host cell raises questions about invasion strategy of *L. monocytogenes;* do multiple bacteria cooperate to increase the likelihood of replication or in contrast, are multiple bacteria cleared more efficiently by host cells. To address this, we tested whether the probability of replication depended on the number of *L. monocytogenes* associated with cells at t0. We could not follow multiple bacteria in the same host cells since it would require much higher (at least one order of magnitude) temporal resolution; instead, based on the live-cell microscopy movies we accurately determined whether the number of bacteria per host cell increases over time or not (Fig. 2c). If *n* denotes the number of bacteria per cell at t0, and *p_0_* is the replication probability when one bacteria is present, then assuming statistical independence the expected probability that at least one bacteria replicates when n is present can be defined as p_n_=1-(1-p_0_)^n^. Cooperativity between bacteria would be associated with increased probabilities of replication (in comparison to the independence model), while increased immune response would be associated with reduced replication probabilities as the number of host-cell associated bacteria increases (Fig. 2d). We found that the observed probabilities exhibited sub-linear increases for up to 7 bacteria per host at t0 (for which at least 10 cells per biological replicate and 50 overall was observed, Fig. 2e). For example, the probability of replication (of at least one bacterium) if two were present at t0 was p_2_=0.29 (±0.05), which increased to p_5_=0.62 (±0.02) when five bacteria were present. We found that the statistical independence model accurately recapitulates the data, with the expected p_0_=0.17 (±0.01). This demonstrates that regardless of the number of bacteria per host, each bacterium has the same probability to establish a replicative invasion, thus acts independently and non-cooperatively.

We noted that the distribution of number of bacteria associated with host cells at t0 exhibits substantial heterogeneity; 33% of host cells were not infected, while some macrophages harboured up to 15 bacteria (Fig. S1a). If the bacterial association was due to a purely random process, the distributions of associated bacteria should follow a one parameter Poisson distribution (Haight 1967). However, the Poisson fit could not capture the data suggesting that a more complex process is involved. Indeed, we found that a negative binomial distribution accurately captures both the increased number of host cells with a very high pathogen number as well as reduced number of those with few or zero bacteria at t0. This suggests that bacterial association with host cells is not purely random, but rather some host cells appear to be more susceptible. For example, in the invasion of non-immune cells, specific ligand/receptor interactions between *L. monocytogenes* and the host are required (Radoshevich and Cossart 2018), which may explain heterogeneity of cell adhesions observed in endothelium (Rengarajan and Theriot 2020). However, invasion of macrophages is a passive process, where *L. monocytogenes* is taken up by host cells through phagocytosis (Flannagan, Cosio, and Grinstein 2009). We found that the number of adherent bacteria was negatively correlated with the number of neighbouring cells (correlation coefficient R^2^= 0.47, p-value <0.01, Fig. S1b), suggesting that adherence is mainly driven by physical accessibility. This suggested that at least in our infection experiments, isolated host cells are more likely to be infected by *L. monocytogenes* than those surrounded by neighbours, probably through increased cell surface available for bacteria to bind.

Overall, these analyses indicate that fate of individual *L. monocytogenes* are independent and non-cooperative in the same host, and the ability to replicate is controlled by behaviour of bacteria.

### Replication probability depends on MOI via phagocytosis

Our data demonstrate that approximately a third of bacteria was able to establish a replicative infection at MOI 0.25 (Fig. 1c). Surprisingly, when the infection was performed at MOI 5 the replication probability was reduced 2-fold (Fig. 3a). Specifically, the probability of replication when one bacterium was present (p_1_) was reduced from 0.32 (±0.06) for MOI 0.25 to 0.17 (±0.01) for MOI 5, while the expected probabilities for multiple bacteria exhibited distinct trends. These data demonstrate that changing the MOI affects the overall replication probability.

**Figure 3.**
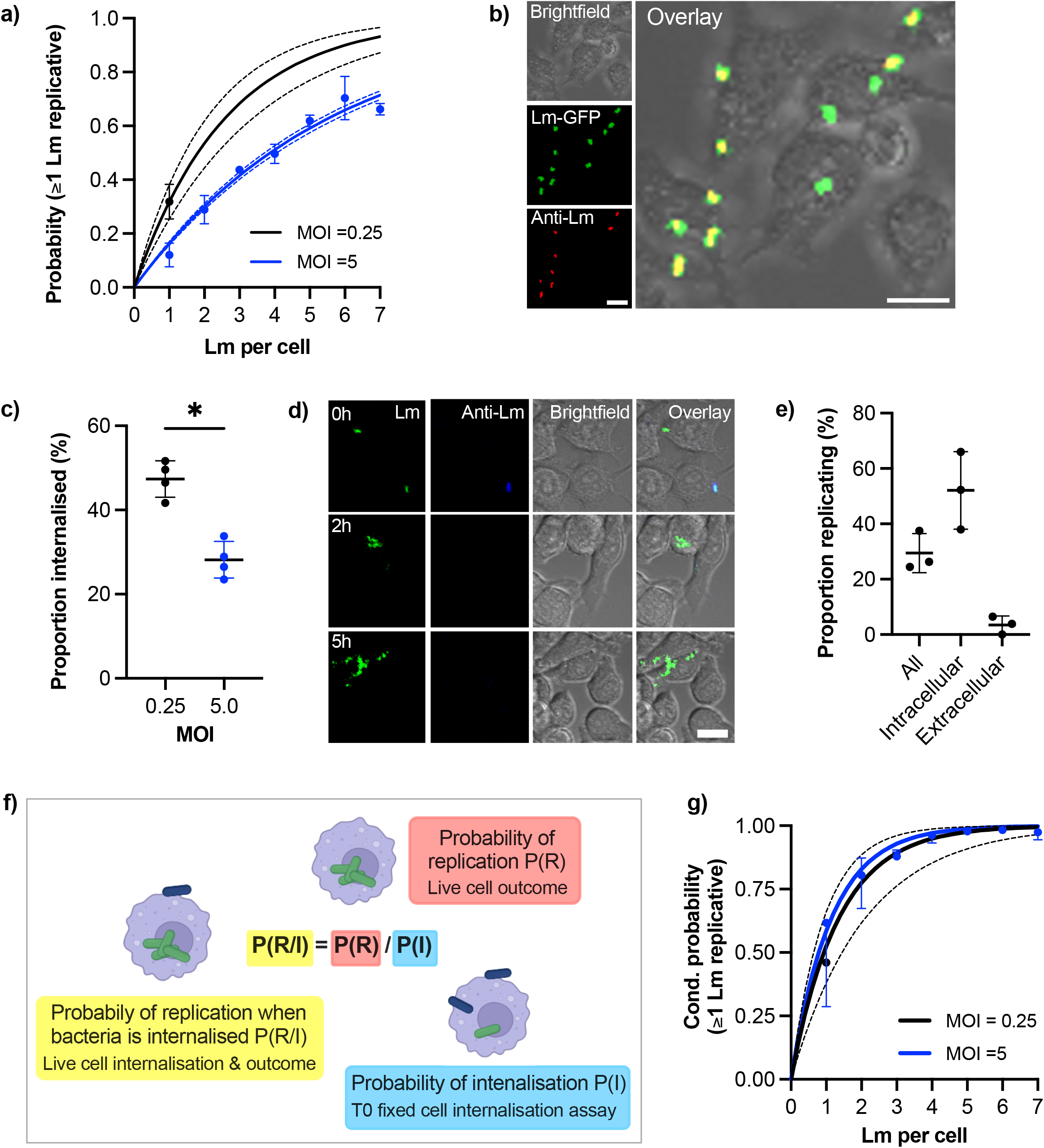
Phagocytosis affects replication probability. a) Probability of replication depends on MOI. Shown is the probability at least one *L. monocytogenes* replicates as a function of number of bacteria per host cell at t0. Solid black line is the predicted probability (with SDs in broken lines) for MOI 0.25, calculated for multiple invasions per host cell given the replication probability p_0_=0.32 (±0.06) for one bacterium per cell (black circle, from Fig. 1c). Similarly, in blue the observed (circles denoting mean with SDs) and expected (solid line with broken line SDs) probabilities for MOI 5 (from Fig. 2e). b) Representative images from internalisation assay showing RAW 264.7 macrophages (brightfield) infected with *Lm*-GFP (green) at MOI 5 fixed at t0 and stained with anti-Lm 594 (red). Scale bar 10 μM. c) Rate of phagocytosis depends on the MOI. The proportion of intracellular *Lm*-GFP at t0 for MOI 0.25 and 5.0, as obtained from assay depicted in b. Individual data points (circles) from four biological replicates with solid lines indicating mean and SD. Statistical significance (* = p-value <0.05) assessed using Mann-Whitney two sample test. d) Representative images of live-cell infection with *Lm*-GFP internalisation staining and infection outcome tracking. Shown are *Lm*-GFP (green), anti-Lm (blue) and RAW 264.7 (brightfield) infection at MOI 0.25, at indicated times. Scale bar 10 μM. e) Proportion of replicating bacteria based on the internalisation status as depicted in d. Shown is proportion replicating of the total *Lm*-GFP associated with a host cell at t0 (all), proportion of the internalised at t0 (intracellular) or proportion of the bacteria that is associated but are not internalised at t0 (extracellular), evaluated at 5 h. RAW 264.7 macrophages infected with *Lm*-GFP at MOI 0.25 using anti-Lm 421 antibody to mark extracellular bacteria at t0. Shown are individual data points (circles) from three replicates with solid lines indicating mean and SD. f) Schematic representation of the conditional probability of replication based on the probability of replication being adjusted to account for the contribution of probability of internalisation. g) Phagocytosis rate explains MOI-specific replication probabilities. Shown are conditional probabilities (of replication given internalisation as described in f) of at least one bacterium replicating as a function of number of bacteria per host cell at t0. Black line indicates the expected probability (mean with SDs) calculated for multiple invasions per host cell given the replication probability p_0_=0.52 ±0.14 (black circle) of internalised bacteria for MOI 0.25 (from Fig. 3e). Blue circles denote conditional probabilities (mean and SDs, based on three replicate experiments) for MOI 5, calculated from data in Fig. 2a based on the proportion of internalised bacteria in Fig. S2. Solid blue line depicts expected conditional probabilities assuming statistical independence and a single overall internalisation rate (0.28 ±0.04 from c) based on the probabilities in Fig. 2e.

In order to replicate bacteria must enter the host cell through phagocytosis and escape to the cytoplasm (Cossart 2011). To test how MOI affects replication probability, we used anti-*Lm* antibody to distinguish bacteria that were internalised from those that were adhering to the cell surface at t0 (Fig. 3b). The proportions of internalized bacteria at t0 was significantly reduced in MOI 5 (28.12 ±4.4%) compared to that of MOI 0.25 (47.4 ±4.3%) demonstrating that higher MOI reduced the rate of phagocytosis (Fig. 3c). By combining live-cell imaging of *Lm*-GFP with anti-Lm extracellular staining (Fig. 3d, video 3), we determined that for MOI 0.25 52% (±14%) of internalized bacteria replicated, while only 3.5% (±1.9%) of the bacteria that were extracellular at t0 replicated (Fig. 3e). The latter likely correspond to bacteria that are in the process of internalisation at t0. We then introduced conditional probabilities to characterise the two-step internalisation/replication process, such that conditional probability of replication given that bacteria is internalised P(R/I)=P(R)/P(I) is the ratio of the overall replication probability P(R) and the probability of internalisation P(I) (Fig. 3f). At MOI 0.25 the expected conditional probability of replication given that one bacteria is internalised P(R/I)=0.62, based on the measured overall replication probability P(R)=0.29 (Fig. 3e) and the probability of internalisation P(I)=0.47 (Fig. 3c). Therefore, the expected probabilities obtained using fixed cell internalisation assay and observed conditional proportions based on dual live-cell staining assay are in the good agreement.

Having confirmed the conditional probability model, we next examined whether the change of the internalisation rate might explain apparent differences in probability of replication observed at different MOIs. While the Fig. 3c captured the overall internalisation rate for MOI 5, we additionally examined the proportion of internalised bacteria as a function of number of bacteria at t0 (Fig. S2). We found a small but significant linear increase from 26.4 (±1.3%) when one bacterium is present up to 41.6 (±5.7%) when seven bacteria were present (R^2^ =0.27, p-value 0.01), suggesting that increased number of bacteria may increase rate of phagocytosis or, according to our previous finding, isolated cells exhibit higher rate of phagocytosis (Fig. S1b). Nevertheless, the conditional probabilities that at least one bacterium replicates calculated for MOI 0.25 and MOI 0.5 based on the associated internalisation rates (see Materials and Methods for derivations) followed almost identical trends (Fig. 3g, black solid line vs. blue circles). The conditional probability calculated for MOI 5, assuming a constant (overall) internalisation rate (from Fig. 3c, blue line) was also in a good agreement with conditional probabilities for MOI 0.25. Therefore, these analyses demonstrate that the changes of replication probability in response to MOI is controlled through the rate of the phagocytosis.

### Population-level secreted LLO levels regulate single cell replication probability

To mechanistically understand how the replication probability depends on the different number of bacteria in the environment we devised a dual colour experiment where *Lm*-GFP (green Lm) equivalent of MOI 0.25 was supplemented with *Lm*-dsRed (red Lm), such that the overall MOI was maintained at 5 (Fig. 4a). We found that only addition of live, but not PFA fixed *Lm*-dsRed significantly reduced the rate of replication, in comparison to the control *Lm*-GFP at MOI 0.25 (Fig. 4b). This suggests a role for a secreted factor produced by bacteria, which is consistent with the soluble pore-forming toxin LLO (Hamon et al. 2012). Indeed, we found when WT *Lm*-GFP were supplemented with *Δhly Lm*-dsRed, unable to produce LLO, the replication probability was not affected at an MOI of 5 (Fig. 4c). Replication probability was similarly affected by live WT but not *Δhly L. monocytogenes* upon invasion of BMDMs (Fig. 4d). LLO is known to play multiple roles during invasion, including activation of host immune responses (Zhang et al. 2019; Lam et al. 2011; Kayal and Charbit 2006; Hamon et al. 2012), we therefore tested whether recombinant LLO alone can inhibit *L. monocytogenes* replication and internalisation. Indeed, we found that incubation with recombinant LLO significantly reduced replication at MOI=0.25 (Fig. 4e), which in BMDMs resulted in almost complete inhibition with <1% bacteria able to establish replicative invasions (Fig. 4f). Consistent with a role for phagocytosis, we observed limited changes of internalisation of *Δhly* strain at MOI 0.25 vs 5, in contrast to the WT bacteria (Fig. 4g). Finally, we showed that treatment with recombinant LLO also significantly reduced the internalisation of bacteria (Fig. 4h). Overall, these data demonstrate that the amount of LLO in the environment surrounding the host macrophages regulates the overall *L. monocytogenes* replication.

**Figure 4.**
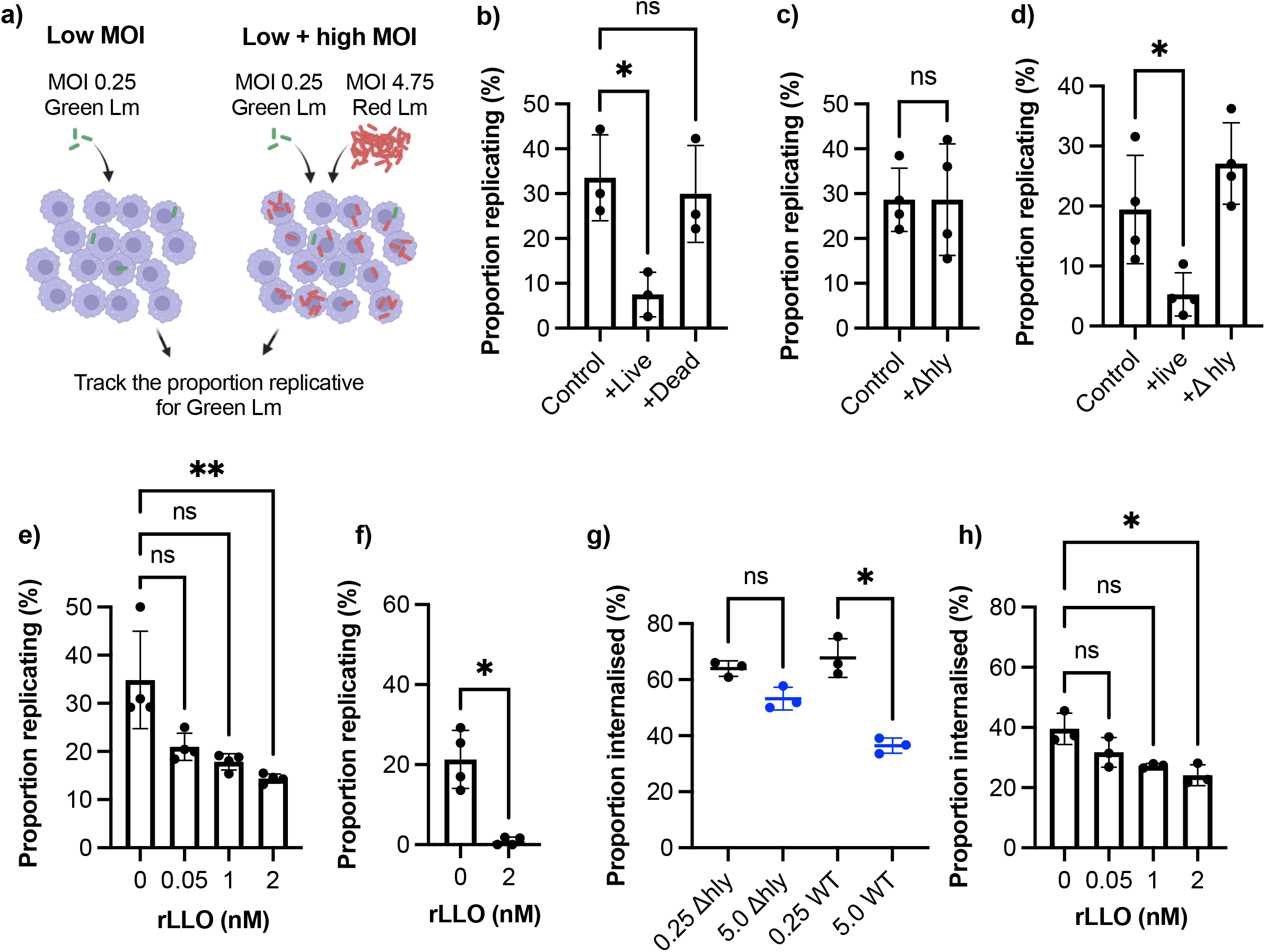
PrfA virulence factors controls infection outcomes at the population-level. a) Schematic representation of experimental set up for infections with *Lm*-GFP and *Lm*-mCherry strains at combined MOI 5 vs *Lm*-GFP MOI 0.25 control. b) Proportion of *Lm*-GFP MOI 0.25 replicating upon infection of RAW 264.7 macrophages for *Lm*-GFP only (control), or for *Lm*-GFP when live (+live) or PFA fixed (+dead) *Lm*-mCherry added for combined MOI 5. Triplicate data (circles) with mean and SD (solid lines). Statistical significance assessed using Kruskal-Wallis ANOVA with Dunn’s correction for multiple comparisons (ns = non-significant, * = p-value<0.05). c) Proportion of *Lm*-GFP (MOI 0.25) replicating upon infection of RAW 264.7 macrophages for *Lm*-GFP only (control), or for *Lm*-GFP when *Δhly Lm*-mCherry (+Δ*hly*) added for combined MOI 5. Replicate data from four experiments (circles) with mean and SD (solid lines). Statistical significance assessed with Mann-Whitney rank test (ns = non-significant). d) Proportion of *Lm*-GFP replicating upon infection of BMDMs at MOI (0.25) (control), or when live WT *Lm*-mCherry (+live) or *Δhly Lm*-mCherry (+Δ*hly*) added for combined MOI 5. Data from four experiments (circles) with mean and SD (solid lines). Statistical significance assessed using Kruskal-Wallis ANOVA with Dunn’s correction for multiple comparisons (ns = non-significant, * = p-value<0.05). e) Proportion of *Lm*-GFP replicating upon infection of RAW 264.7 macrophages at MOI 0.25, with addition recombinant listeriolysin (rLLO). Individual data from four experiments (circles) with mean and SD (solid lines). rLLO at indicated concentrations added with inoculant and removed at t0. Statistical significance assessed using Kruskal-Wallis ANOVA with Dunn’s correction for multiple comparisons (ns = non-significant, ** = p-value<0.01). f) Proportion of *Lm*-GFP replicating upon infection of BMDM at MOI 0.25, incubated with 0 or 2 nM rLLO. Individual data from three experiments (circles) with mean and SD (solid lines). rLLO added with inoculant and removed at t0. Statistical significance assessed with Mann-Whitney rank test (* = p-value<0.05). g) Proportion of WT *Lm*-GFP and *Δhly Lm*-GFP internalised into RAW 264.7 macrophages at MOI 0.25 and 5. Data obtained from internalisation assay using anti-Lm 594 staining of infected cells fixed at t0 as depicted in 3b. Values from triplicate data (circles) with mean and SD (solid lines). Statistical significance assessed using Kruskal-Wallis ANOVA with Dunn’s correction for multiple comparisons (ns = non-significant, * = p-value<0.05). h) Proportion of *Lm*-GFP internalised into RAW 264.7 macrophages at MOI 0.25 in the presence of recombinant LLO (rLLO). Individual replicate data from four experiments (circles) with mean and SD (solid lines), from internalisation assays using anti-Lm 594 staining of infected cells fixed at t0. rLLO concentration indicated on the graph. Statistical significance assessed using Kruskal-Wallis ANOVA with Dunn’s correction for multiple comparisons (ns = non-significant, * = p-value<0.05).

### PrfA activity is necessary but not sufficient to induce replication at the single cell level

Given the role of the PrfA-mediated virulence in the control of the overall replication probability, we wanted to understand whether the PrfA activity also determined the replication at the single cell level. To follow virulence expression in individual cells we developed dual reporter *Lm*strains, in which appropriate chromosomally integrated promoter (*Phly* or *PactA*) drives expression of GFP, in addition to constitutively expressed tagRPF (see Materials and Methods). In agreement with previous analyses (Guldimann et al. 2017), we found that the expression of *Phly*-GFP exhibited substantial heterogeneity in WT *L. monocytogenes* when cultured in tissue culture cell media for up to 1.5h, with only a subset of cells reaching high expression levels (Fig. S3). While a control *ΔprfA* showed no detectable *Phly*-GFP expression, the PrfA* strain, in which PrfA is constitutively activated (Reniere et al. 2015), exhibited substantially elevated fluorescence levels and reduced cell-to-cell variability (coefficient of variation 0.93 vs. 0.62) comparing to that of WT (Fig. S3).

We hypothesised that the ability to establish a replicative invasion was dependent on the level of PrfA activity in individual bacteria. We therefore used confocal microscopy to follow temporal regulation of PrfA activity and fate of individual bacteria upon infection of RAW 264.7 macrophages (Fig. 5a, video 4-5). First, we found that following invasion, representative replicating bacteria tracked with a high temporal resolution (every 5 min) exhibited induction of PrfA activity (Fig. 5b). Specifically, *Phly*-GFP expression rapidly increased and was maintained within the 2 h time window, notably through multiple division events. Similarly, *PactA*-GFP was robustly induced with a delayed kinetics, as previously indicated at the population level (Bubert et al. 1999) and predicted by differences in PrfA-PrfA box binding specificities between *hly* and *actA* promoter regions (Scortti et al. 2007). The robust activation from both promoters was observed through multiple divisions, which suggested an ongoing transcription. In contrast, representative bacteria that did not replicate showed lower PrfA activity, however at least one non-replicative tracked bacterium robustly upregulated *Phly*-GFP expression (depicted in blue in Fig. 5 a and b).

**Figure 5.**
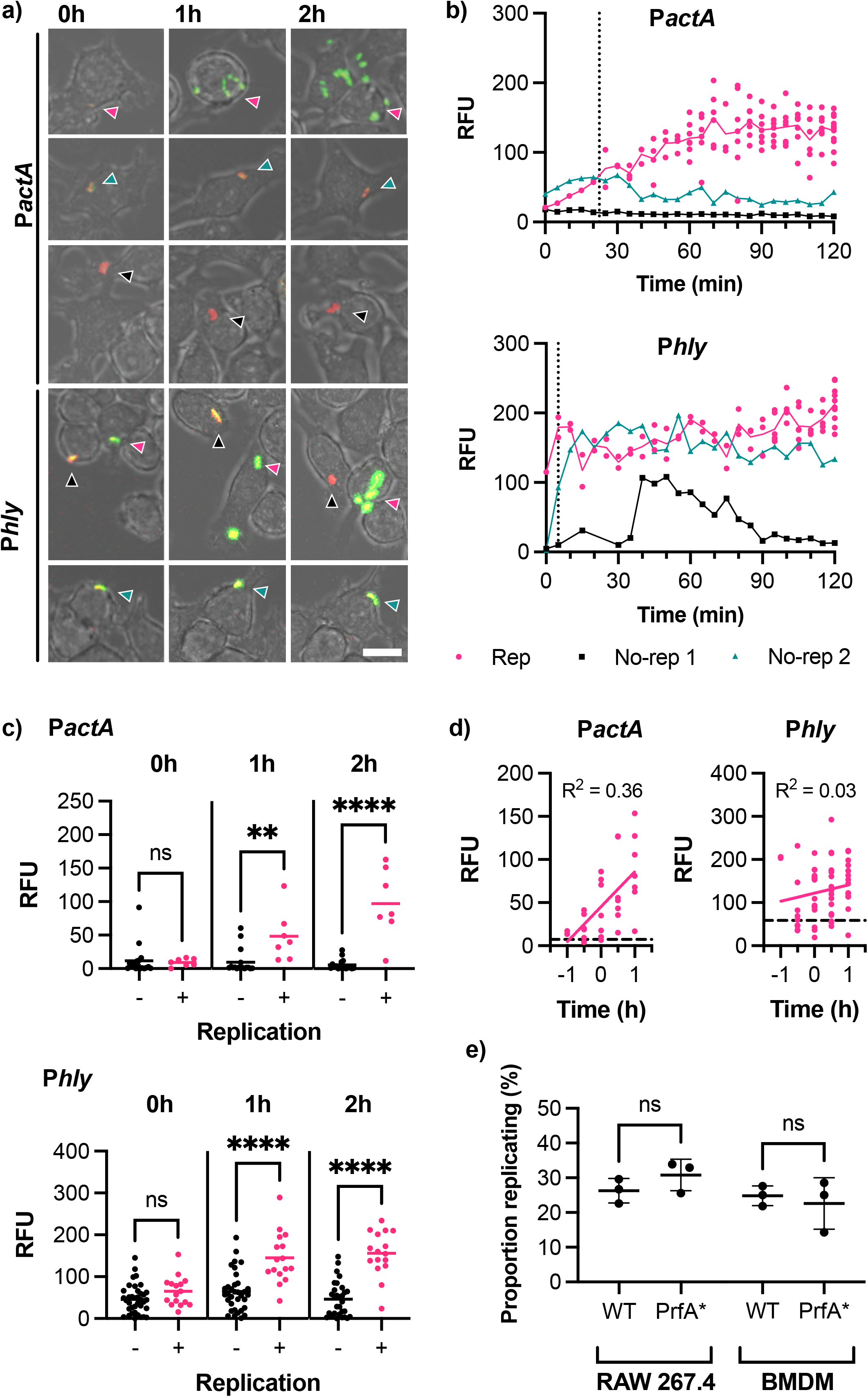
PrfA activity correlates with, but does not determine, infection outcome. a) Representative images of live-cell *Lm*-dsRed P*actA*-GFP (P*actA*) or *Lm*-dsRed P*hly*-GFP (P*hly*) infection of RAW 264.7 macrophages at MOI 0.25. Shown are RAW 264.7 macrophages (brightfield) infected with *Lm*-dsRed P*actA*-GFP or *Lm*-dsRed P*hly*-GFP (red) and expressing GFP under the control of *actA* or *hly* promoter region (green) at 0h, 1h or 2h. Arrows indicate the individual *L. monocytogenes* or replicative foci, for replicative (pink), non-replicative 1 (black) or non-replicative 2 (teal). Scale bar 10 μM. b) Reporter expression trajectories over time for representative individual *Lm*-dsRed *PactA*-GFP (P*actA*) or *Lm*-dsRed P*hly*-GFP (P*hly*) and their daughter cells during infection of RAW 264.7 macrophages at MOI 0.25 from 0-2h. Individual tracked bacteria that were replicative (Rep, pink) or non-replicative (no-rep 1, black; no-rep 2, teal) indicated by circles and correspond to the images in a. GFP intensities measured as relative fluorescence units (RFU) every 5 min, for up to 12 (P*actA*) or 10 (P*hly*) replicative daughter cells. Mean RFU (solid lines) and time of first replication (dotted line) also shown. c) Reporter fluorescence expression for *Lm*-dsRed P*actA*-GFP (P*actA*) or *Lm*-dsRed P*hly*-GFP (P*hly*) cells during infection of RAW 264.7 macrophages at MOI 0.25. Data from 3 replicate experiments for individual non-replicative bacteria (black circles, total 37 for *Phly*, 17 for P*actA*) or representative individual bacteria from all replicative foci (pink circles, total 16 for *Phly*, 7 for P*actA*) and their mean (solid lines) shown for 0, 1 and 2h. GFP intensities measured as relative fluorescence units (RFU). Statistical significance (ns = non-significant, ** = p-value <0.01, **** = p-value <0.0001) assessed using Mann-Whitney rank test. d) Reporter fluorescence expression by time before/after first replication for *Lm*-dsRed P*actA*-GFP (P*actA*) or *Lm*-dsRed P*hly*-GFP (P*hly*) during infection of RAW 264.7 macrophages at MOI 0.25. Data from 3 replicates for representative individual bacteria from all replicative foci (pink circles), simple linear regression for replicative data (pink solid line) and average non-replicative expression (black broken line) shown (with corresponding correlation coefficient R^2^). GFP intensities measured as relative fluorescence units (RFU). e) Proportion of WT or PrfA* *Lm*-GFP replicating upon infection of RAW 264.7 or BMDM primary macrophages at MOI 0.25. Individual data from three experiments (circles) with mean and SD (solid lines). Statistical significance assessed with Mann-Whitney rank test.

To analyse the patterns of PrfA activity more systematically, we quantified *Phly*-GFP and *PactA*-GFP at selected times post invasion. At t0, the levels of *Phly*-GFP and *PactA*-GFP expression were not statistically different between those bacteria that went on to establish a replicative infection and those that did not. However, at 1 and 2 h post invasion, replicative bacteria induced significantly higher *Phly-GFP* or *PactA*-GFP reporter expression, comparing to those bacteria that did not replicate (Fig. 5c). While some non-replicative bacteria exhibited substantial *Phly-GFP* expression over time, corresponding *PactA*-GFP fluorescence always remained at low basal levels. This suggested a statistical relationship between the elevated levels of PrfA required for activation of the *PactA* promoter and the infection outcome. To refine these analyses, we estimated the time to the (first) replication from the time-lapse imaging data, which typically occurred between 5 to 90 mins from t0 (Fig. S4) and used that to normalise temporal PrfA trajectories according to replication time (Fig. 5d). This analysis demonstrated that temporal increases of *PactA*-expression was a very strong indicator for the replicative invasion, highlighted by a high temporal correlation of expression levels for *PactA* (R^2^= 0.36, p-value <0.001) but not *Phly* (R^2^= 0.03, p-value= 0.21).

To test this apparent correlation functionally, and to specifically quantify if enhanced PrfA activation increases replication probability, we analysed infections with PrfA*-*Lm*, in which PrfA shows higher activity compared to a WT strain (Fig. S3). We found no statistical difference in the replication probability of WT or PrfA* strains in RAW 264.7 and BMDMs macrophages (at MOI 0.25), demonstrating that increased PrfA activity *per se* was not sufficient to induce replication. Overall, these analyses demonstrate that in a marked contrast to population level strategy relying on collective PrfA activation and LLO secretion, the level of PrfA activity does not determine whether bacterium is able to replicate or not at the single cell level.

## Discussion

Interactions between host and pathogen at the single cell level are inherently heterogeneous leading to different infection outcomes. Here we use time-lapse confocal microscopy to follow with a single bacterium resolution the fate of an important food-borne pathogen *L. monocytogenes* upon the invasion of innate immune macrophages, a key event controlling the overall infection. We demonstrate that infection of macrophages results in heterogeneous outcomes, where only a fraction of single-cell host pathogen interactions leads to intracellular replication and spread of bacteria, while many bacteria are cleared or remain in a non-replicative state in the host (Fig. 1). In our datasets ~30% of bacteria were able to establish replicative infection, both in RAW 264.7 line as well as primary murine BMDMs. Successful replication of *L. monocytogenes* is a muti-step process requiring host cell entry and phagosome escape (Radoshevich and Cossart 2018); our data demonstrate that once internalised by a host cell, approximately 50% of individual bacteria can establish a replicative infection, regardless of the level of infection (i.e., low or high MOI, Figs. 2 and 3). In addition, approximately 25% of internalised bacteria persist in the host in a non-replicative state for at least 5 h. Our data demonstrate that at least 30% of these non-replicative bacteria associate with actin, an established marker for cytoplasmic localisation (Cossart 2011), suggesting that they are in the cytoplasm, but do not replicate. Previous work showed that during prolonged invasion the intracellular *L. monocytogenes* may switch between replicative and persistent non-replicative state in the non-phagocytic human cells (Kortebi et al. 2017). The non-replicative state coincides with decline of ActA expression and incorporation into late endosomal/lysosomal vacuoles. Our imaging experiment provide no evidence for the reactivation of the persistent cells (within the 5 h time window), instead the replication occurs as quickly as 30 mins after addition of inoculum, with almost all cells establishing replication within 2 h window post infection.

Pathogens often cooperate to overcome host cell defences (Diard et al. 2013). For example, cooperativity between *Salmonella* allows non-invasive strains to enter host cells (Lorkowski et al. 2014; Kazmierczak, Mostov, and Engel 2001; Misselwitz et al. 2012; Ginocchio, Pace, and Galán 1992). Once in the host, cooperativity among bacterial effector protein enable suppression of the immune defences by targeting multiple signalling responses (de Jong and Alto 2018). In turn, host cells use collective behaviour, including quorum-like activation of their signalling responses, to enhance immune responses (Muldoon et al. 2020) for better pathogen control (Boechat et al. 2001). Using dual colour experiments we found that the invasion strategies of individual *L. monocytogenes* in the same host cell are non-cooperative (Fig 2). Specifically, when multiple bacteria invade the same host cell all act independently with the same probability of replication. Importantly, while not providing any apparent advantage, the presence of multiple bacteria in the same host cell virtually assures a certain replication and subsequent rapid intracellular proliferation of *L. monocytogenes*. For example, the probability of replication for >3 bacteria per host exceeds 90%. Paradoxically, we found that higher MOI resulted in ~2-fold reduction of the replication probability. We demonstrate that this is regulated through the phagocytosis in the host, which can alter the rate of *L. monocytogenes* uptake (Fig. 3). We demonstrate that this is due to expression of PrfA-mediated LLO, a pore forming toxin, which is sufficient to inhibit phagocytosis and subsequent replication in LLO treated cells (Fig. 4). At the single cell level, we show that the major PrfA regulon is necessary, but not sufficient for intracellular replication (Fig. 5). We demonstrate that while bacteria exhibit substantial heterogeneity of PrfA activity, and replicative bacteria maintain high PrfA activity (to drive ActA reporter expression), increased PrfA activity does not lead to more replication events. Which pathway in *L. monocytogenes* controls successful replication remains unclear, but perhaps one candidate is the DNA uptake competence (Com) system, which is required for phagosome escape and exhibits expression variability and is regulated independently of

PrfA through prophage partial induction (Rabinovich et al. 2012; Pasechnek et al. 2020). However, our data show that while single bacteria act non-cooperatively, bacterial populations use cooperative virulence expression to manipulate host responses. We suggest that reduced phagocytosis *in vivo* might assure successful replication while simultaneously increasing the likelihood of systemic dissemination through the blood stream, and uptake by non-phagocytic cells (Dramsi and Cossart 2003) to promote immune evasion.

LLO is a pore-forming toxin, pH-dependent member of the cholesterol-dependent cytolysins which binds cholesterol present in the host cell membrane (Hamon et al. 2012). It is necessary for the vacuolar escape of *L. monocytogenes*, but plays many other roles, including control of autophagy and mitophagy (Zhang et al. 2019), and suppression of ROS production (Lam et al. 2011), but also activates host signalling responses (Kayal and Charbit 2006). Previous work suggests that the formation of LLO pores at the cell membrane has been shown to induce *L. monocytogenes* internalisation into non-phagocytic cells (Dramsi and Cossart 2003; Vadia et al. 2011). Our data demonstrate that in macrophages, the rate of phagocytosis decreased upon LLO exposure, while the effect on overall ability to establish replicative invasion, especially in primary macrophages is substantial. Phagocytosis was previously shown to be regulated, in part through p38 mitogen activated protein kinase, in response to TRL2-dependent Gram-positive *Staphylococcus aureus* (Doyle et al. 2004; Blander and Medzhitov 2004) and *L. monocytogenes* (Shen et al. 2010). Phagocytosis of *L. monocytogenes* has also been linked to the expression of the inhibitory receptor T-cell immunoglobin mucin-3 (Tim-3), an immune checkpoint inhibitor (Wolf, Anderson, and Kuchroo 2020). Tim-3 inhibits the rate of phagocytosis by inhibiting expression of the CD36 scavenger receptor (Wang et al. 2017), which is involved in phagocytosis of Gram positive bacteria (Baranova et al. 2008). Tim-3 itself and its ligand Galectin-9 both have been shown to be upregulated by infection (Jayaraman et al. 2010). In addition, LLO (and other cholesterol cytolysins) have been shown to bind the mannose receptor (MCR1), while blocking MRC1 resulted in reduced uptake and intracellular survival of *Streptococcus pneumoniae* (Subramanian et al. 2020). It is currently unclear, whether physiological levels of LLO may indeed be sufficient to alter the expression of these receptor system and thus alter phagocytosis. Nevertheless, these data suggest that control of phagocytosis via cholesterol cytolysins might represent an important invasion strategy for bacterial pathogens. In agreement, pneumonolysin, which is structurally similar to LLO, was also shown to inhibit phagocytosis of *S. pneumoniae* in neutrophils (Ullah, Ritchie, and Evans 2017). In turn, the control of phagocytosis and phagosome maturation remains an important host defence strategy (Drevets, Leenen, and Campbell 1996; Calame, Mueller-Ortiz, and Wetsel 2016) (Kernbauer et al. 2012; Dalton et al. 1993; Buchmeier and Schreiber 1985), highlighting the critical role of phagocytosis in host pathogen interactions.

Overall our analyses reveal new insight into distinct single cell and population-level strategies of *L. monocytogenes* upon invasion of innate immune macrophages. We demonstrate that while inside the host cells, individual bacteria act independently and non-cooperatively, the overall bacterial population control outcomes of single cell host interactions through collective PrfA signalling.

## Conflict of Interest

The authors declare that the research was conducted in the absence of any commercial or financial relationships that could be construed as a potential conflict of interest.

## Author contributions

JM developed bacterial strains for imaging, collected, and analysed data. JB provided preliminary data for the project. MG and EL assisted with strain development. LF, CN and DS assisted with imaging analyses. IR and PP provided supervision and conceptualisation, and with assistance of JM wrote the manuscript. All authors read and approved the final manuscript.

## Funding

This work was supported by BBSRC (BB/R007691/1).

## Acknowledgments

BMDMs were a kind gift from Dominik Ruckerl and Jack Green. Plasmids pPL2-mCherry, pAD3-PactA-GFP (BUG2793) and pPL2-Phly-GFP (pCG8) were gifts from John-Demian Sauer, Pascale Cossart and Claudia Guldimann respectively.

## Data Availability Statement

Tabularized manuscript data will be provided via public repositories upon publication.

## Materials and Methods

### Bacterial strains culture conditions

*L. monocytogenes* EGDe:InlA^m^ (Wollert et al. 2007) was used as the wild type (WT), with all mutations generated in this background. *L. monocytogenes* was grown in tryptone soya broth (TSB) unless otherwise stated, when needed antibiotics were added at final concentrations of: chloramphenicol (Cm) 7 μg ml^-1^ and erythromycin (Em) 5 μg ml^-1^. *Escherichia coli* DH5α was used for cloning and grown in Luria-Bertani broth (LB), when needed antibiotics were added at final concentrations of: chloramphenicol (Cm) 35 μg ml^-1^ and erythromycin (Em) 150 μg ml^-1^.

Plasmids (Table 1) were electroporated into *L. monocytogenes* to generate fluorescently tagged and fluorescent reporter strains of *L. monocytogenes* described in the same table. Chromosomal integration of integrative plasmids was confirmed by PCR as described previously (Lauer et al. 2002). Correct fluorescence of strains was confirmed by microscopy.

**Table 1.** Plasmids and strains used in this study

| Plasmid, strain or primer name | Description | Antibiotic resistance | Reference |
| --- | --- | --- | --- |
| **Plasmids** |  |  |  |
| pAD<sub>1</sub>-cGFP | Integrative plasmid with constitutive GFP expression | Cm | (Balestrino et al. 2010) |
| pAD<sub>3</sub>-PactA-GFP | Integrative plasmid expressing GFP under control of *PactA* | Cm | (Balestrino et al. 2010) |
| pCG8 | Integrative plasmid expressing codon optimised GFP under control of *Phly* | Cm | (Guldimann et al. 2017) |
| pJEBAN6 | Plasmid expressing constitutive DsRedExpress | Em | (Andersen et al. 2006) |
| pPL2-mCherry | Integrative plasmid expressing constitutive codon optimised mCherry | Cm | (Vincent et al. 2016) |
| **Bacterial strains** |  |  |  |
| EGDe:InIA<sup>m</sup> | EGDe strain with murinized InIA protein | - | (Wollert et al. 2007) |
| $\Delta hly$ *Lm* | EGDe:InIA<sup>m</sup> *hly* deletion mutant | - | (Wang et al. 2015) |
| $\Delta prfA$ *Lm* | EGDe:InIA<sup>m</sup> *prfA* deletion mutant | - | This study |
| PrfA* *Lm* | EGDe:InIA<sup>m</sup> PrfA* mutant | - | This study |
| *Lm*-GFP | EGDe:InIA<sup>m</sup> with integrated pAD<sub>1</sub>-cGFP | Cm | This study |
| *Lm*-DsRed | EGDe:InIA<sup>m</sup> with pJEBAN6 | Em | This study |
| *Lm*-mCherry | EGDe:InIA<sup>m</sup> with integrated pPL2-mCherry | Cm | This study |
| $\Delta hly$ *Lm*-GFP | $\Delta hly$ *Lm* with integrated pAD<sub>1</sub>-cGFP | Cm | This study |
| $\Delta hly$ *Lm*-mCherry | $\Delta hly$ *Lm* with integrated pPL2-mCherry | Cm | This study |
| *Lm*-dsRed *PactA*-GFP | EGDe:InIA<sup>m</sup> with integrated pAD<sub>3</sub>-*PactA*-GFP and pJEBAN6 | Cm, Em | This study |
| *Lm*-dsRed *Phly*-GFP | EGDe:InIA<sup>m</sup> with integrated pCG8 and pJEBAN6 | Cm, Em | This study |
| PrfA* *Lm*-GFP | PrfA* *Lm* with integrated pAD<sub>1</sub>-cGFP | Cm | This study |
| $\Delta prfA$ *Lm*-dsRed *Phly*-GFP | $\Delta prfA$ *Lm* with integrated pAD<sub>3</sub>-*PactA*-GFP and pJEBAN6 | Cm, Em | This study |
| PrfA* *Lm*-dsRed *Phly*-GFP | PrfA* *Lm* with integrated pAD<sub>3</sub>-*PactA*-GFP and pJEBAN6 | Cm, Em | This study |

*L. monocytogenes* PrfA* and *ΔprfA* mutants were constructed using the temperature sensitive shuttle plasmid pAUL-A as described previously (Wang et al. 2015).

### Cell culture

RAW 264.7 macrophages were maintained in Dulbecco’s modified Eagle medium (DMEM) supplemented 10% (v/v) fetal bovine serum (FBS) and 1% (v/v) 100x MEM non-essential amino acid solution (NEAA) at 37 °C 5% (v/v) CO_2_. Bone marrow derived macrophages (BMDMs) were generated from C57BL/6 female mice using L929-conditioned media, once differentiated BMDMs were maintained for up to 3 days in DMEM supplemented with 10% (v/v) FBS at 37 °C 5% (v/v) CO_2_.

### Live-cell microscopy infection assays

RAW 264.7 macrophages or BMDMs were seeded in 35 mm TC treated imaging dishes (Cellview Greiner) at 3.5×10^5^ and 7×10^6^ cells ml^-1^ respectively and incubated overnight. *L. monocytogenes* mid-log (OD_600_ 0.45-0.6) aliquots stored at −80 °C in PBS glycerol (15% v/v) were used for infections. Cells were infected with *L. monocytogenes* at a MOI of 0.25 in prewarmed media for 45 min and washed three times prior to the addition of 10 μg ml^-1^ gentamicin media (Fig. 1a). For assays with recombinant listeriolysin (rLLO, Abcam), rLLO was added to cells with the *L.monocytogenes* at a final concentration of 0.05-2 nM. Infections were immediately imaged by live-cell time-lapse microscopy using a Zeiss LSM710, Zeiss LSM780 or Zeiss LSM880 microscope. Data was visualised using the Zeiss Zen Black software.

### Actin staining and internalisation assay

For internalisation assay and actin staining assay infections were performed as described above but fixed at 0 h and 5 h respectively with 4% (w/v) PFA PBS for 30 min at room temperature. For the internalisation assay, extracellular *L. monocytogenes* were stained with polyclonal rabbit anti-listeria (anti-Lm) antibody (Abcam, ab35132), and washed 3 times before secondary antibody staining with anti-rabbit IgG 594 (Sigma-Aldrich). For the live-cell internalisation and infection outcome assay, anti-Lm was added to live cells in pre-warmed media for 30 s, washed 3 times before secondary antibody staining with Brilliant violet 421 donkey anti-rabbit antibody (Biolegend) in pre-warmed media for 30 s, and washed 3 times before imaging.

For the actin staining assay fixed cells were permeabilised 0.1% triton X-100 (v/v) PBS for 4 min and washed 3 times with PBS. As permeabilization sometimes affected *Lm*-GFP signal intensity, anti-Lm staining was then performed as described above, but with anti-rabbit IgG 488 (Biolegend) secondary antibody. Alexa fluor 594 Phalliodin (ThermoFisher) was used to stain actin.

### Analysis of imaging data

To identify infection outcomes in time-lapse microscopy data, individual *L. monocytogenes* visible at t0 were visually tracked in Zen Black software and recorded as bacterial replication, no-replication or bacteria disappear (Fig. 1b). Correlations between actin or anti-Lm staining were manually assessed in Zen Black.

For tracked *L. monocytogenes* GFP reporter expression (*Phly* or P*actA*) during infection, individual *L. monocytogenes* were highlighted as regions of interest for selected time points in FIJI (Schindelin et al. 2012), and relative fluorescence intensities (RFU) exported for downstream analysis.

For *L. monocytogenes* P*hly*-GFP reporter expression in media at selected timepoints, automated analysis of exported tif images was performed in CellProfiler (McQuin et al. 2018). Brightfield images were used to segment images and identify bacterial cell outlines, relative fluorescence intensities for individual bacteria were then exported for downstream analysis.

### Analysis of replication probabilities

In general, the conditional probability that bacteria replicates (R) given that it is internalised (I); P(R/I), can be expressed as the ratio of the overall replication probability P(R) and the internalisation probability P(I); such that P(I)=P(R)/P(I). Then P_n_(I) = P_n_(R)/P_n_(I) denotes respective probabilities for n bacteria adhering to the host cell at t0. The expected conditional probability that at least one bacteria replicates is given by p_n_(R/I)=1-(1-P_n_(R/I))^n^, where in general P_n_(R/I) may depend on n. However, under the statistical independence model (Fig. 2d), these relationships are equivalent to p_n_(R/I)=1-(1-p_0_(R/I))^n^ where n is the number of adherent bacteria and p_0_(R/I)=I(R/I)= P_1_(R)/P_1_(I) is the probability of replication if one bacteria is present. For MOI 0.25, p_0_(R/I) was measured directly using live-cell microscopy with additional staining (Fig. 3d), and subsequently used to calculate expected probabilities for n>1 (Fig. 3g, blue curve). For MOI 5, we used a previously fitted p_0_=0.164 (Fig. 2e) such that p_0_=P_n_(R) and p_n_(R/I)=1-(1-p_0_/P_n_(I))^n^. Then the probability of internalisation P_n_(I) was either measured for each n (Fig. S2) or a single average rate was used (as in Fig. 3c).

### Statistical analyses

Statistical analysis was performed using GraphPad Prism 8 software (version 8.4.2). The D’Agostino-Pearson test was applied to test for normal (Gaussian) distribution of acquired data. Two-sample comparison was conducted using non-parametric Mann Whitney test, for analyses of variance Kruskal-Wallis ANOVA with Dunn’s multiple comparisons test was performed. Simple linear regression and Pearson’s correlation coefficient R^2^ was used to test association between two selected variables.

## Supplementary figures

**Figure S1. Distribution of bacteria adhering to host cells at MOI 5.**

a) Distribution of bacteria per cell at t0 when RAW 264.7 macrophages simultaneously infected with *Lm*-dsRed and *Lm*-GFP at combined MOI 5 (at 1:1 ratio). Plot shows mean frequency of distribution for experimental data (black) from 3 replicates, and expected data assuming a one parameter Poisson distribution (pink) or negative binomial distribution (teal).

b) Relationship between mean frequency of bacteria per cell at t0 and number of neighbouring cells when RAW 264.7 macrophages simultaneously infected with *Lm*-dsRed and *Lm*-GFP at combined MOI 5 (at 1:1 ratio). Data from 2 replicates (black circles) and simple linear regression (black line, R^2^ = 0.47, p-value = 0.007).

**Figure S2. Analysis of *L. monocytogenes* internalisation at MOI 5**. Relationship between proportion of internalised *Lm*-GFP and number of *Lm*-GFP associated with the cell at t0 when RAW 264.7 macrophages infected at MOI 5. Individual data from four replicates (circles) and simple linear regression (solid line, R^2^ = 0.27, p-value = 0.01) shown. Data obtained from internalisation assay using anti-Lm 594 staining of infected cells fixed at t0 as depicted in 3b.

**Figure S3. PrfA operon reporter expression at the single cell level.**

a) Representative images of live-cell WT, *ΔprfA* or PrfA*\* Lm*-dsRed P*hly*-GFP incubated in DMEM over time. Shown are *Lm* cells (brightfield) and expression of GFP under the control of the *hly* promoter region (GFP) at 0.5, 1.5 or 2.5h after addition of *Lm*. Scale bar 5 μM.

b) GFP fluorescence expression from the *hly* promoter over time. Data from 3 replicate experiments (minimum 137 total individual cells per condition) for WT (pink), *ΔprfA* (black) or PrfA*\** (teal) *Lm*-dsRed P*hly*-GFP incubated in DMEM. Automated cell identification and GFP intensity measured as relative fluorescence units (RFU) in Cell Profiler at 0.5, 1.5 or 2.5h after addition of *L. monocytogenes* to media. Statistical significance (ns = non-significant, **** = p-value <0.0001) assessed using Kruskal-Wallis ANOVA with Dunn’s correction for multiple comparisons.

**Figure S4. Distribution of *L. monocytogenes* replication times post invasion.** Shown is the time to first replication of *Lm*-dsRed P*actA*-GFP and *Lm*-dsRed P*hly*-GFP during infection of RAW 264.7 macrophages at MOI 0.25. Data from 3 replicates and 22 total replicative *L. monocytogenes*.

## Videos

V1) Representative live-cell microscopy of RAW 264.7 macrophages (brightfield) infected with *Lm*-GFP (green) from times 0-10 h post gentamycin treatment. Scale 20μM.

V2) Representative live-cell microscopy of RAW 264.7 macrophages (brightfield) simultaneously infected with *Lm*-dsRed (red) and *Lm*-GFP (green) at combined MOI 5 (at 1:1 ratio) from times 0-5 h post gentamycin treatment. Scale bar 10 μM.

V3) Representative live-cell microscopy of RAW 264.7 macrophages (brightfield) infected with *Lm*-GFP (green) with anti-Lm 421 internalisation staining (magenta) from times 0-5 h post gentamycin treatment. Scale bar 10 μM.

V4) Representative images of live-cell *Lm*-dsRed P*actA*-GFP (red) infection of RAW 264.7 macrophages (brightfield) at MOI 0.25 from times 0-5 h post gentamycin treatment. GFP expression is driven from the *actA* promoter region (green). Scale bar 10 μM.

V5) Representative images of live-cell *Lm*-dsRed P*hly*-GFP (red) infection of RAW 264.7 macrophages (brightfield) at MOI 0.25 from times 0-5 h post gentamycin treatment. GFP expression is driven from the *hly* promoter region (green). Scale bar 10 μM.

## Notes

### Competing Interest Statement

The authors have declared no competing interest.

