## Supplementary figures and images for "Single-cell imaging reveals non-cooperative and cooperative infection strategies of *Listeria monocytogenes* in macrophages"

### Figure S1

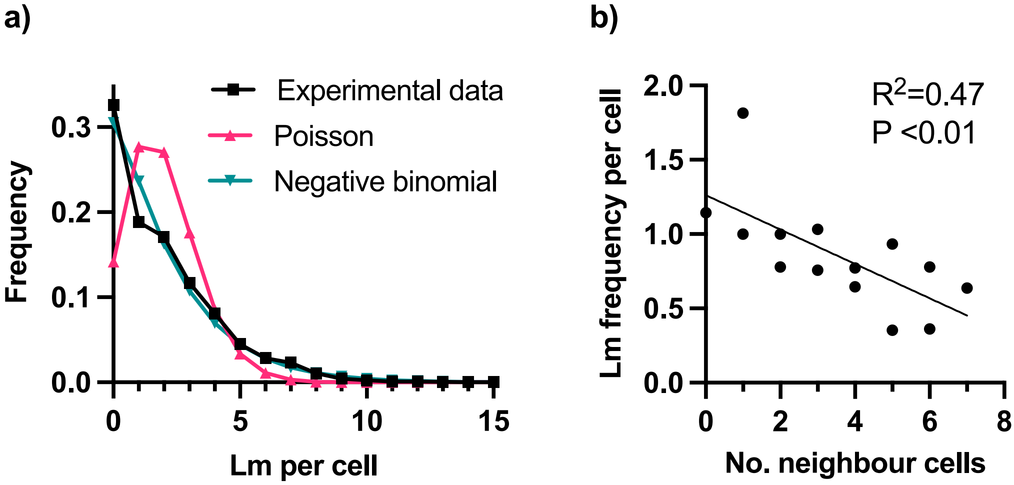

### Figure S2

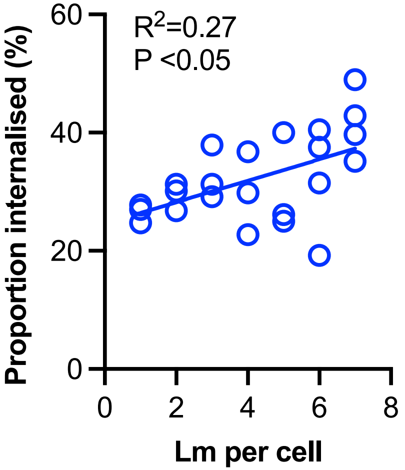

### Figure S3

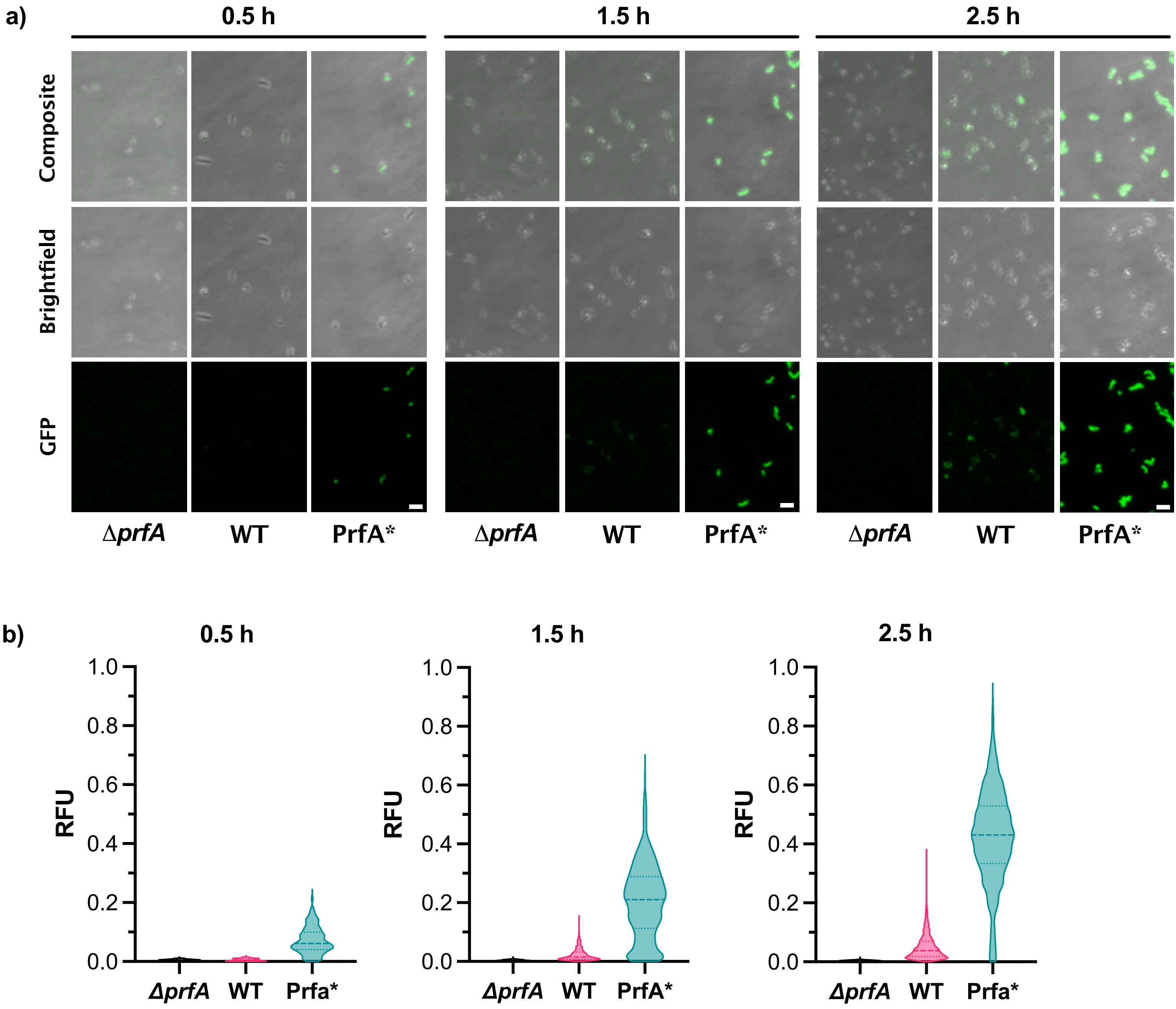

### Figure S4

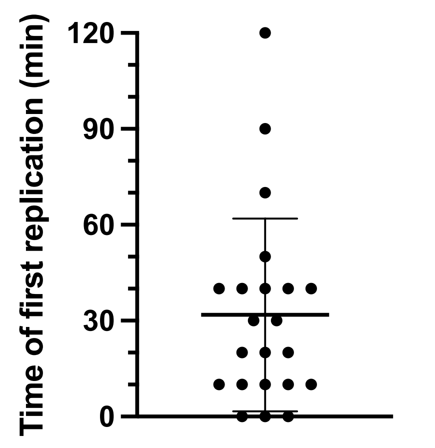
